# The Autism Risk Factor CHD8 Is a Chromatin Activator in Human Neurons and Functionally Dependent on the ERK-MAPK Pathway Effector ELK1

**DOI:** 10.1101/2020.11.10.377010

**Authors:** Bahareh Haddad Derafshi, Tamas Danko, Soham Chanda, Pedro Batista, Ulrike Litzenburger, Qian Yi Lee, Yi Han Ng, Anu Sebin, Howard Y. Chang, Thomas C. Südhof, Marius Wernig

## Abstract

The chromodomain helicase DNA-binding protein CHD8 is among the most frequently found de-novo mutations in autism spectrum disorder (ASD)^1–4^. Despite its prominent disease involvement, little is known about its molecular function in the human brain. CHD8 is believed to be a chromatin regulator, but mechanisms for its genomic targeting is also unclear. To elucidate the role of CHD8 in human neurons, we generated conditional loss-of-function alleles in pluripotent stem cells. Chromatin accessibility and transcriptional profiling showed that *CHD8* is a potent chromatin opener and transcriptional activator of its direct neuronal targets, including a distinct group of ASD genes. We found the chromatin targeting of CHD8 to be highly context-dependent. In human neurons, CHD8 was preferentially bound at promoter sequences which were significantly enriched in ETS motifs. Indeed, the chromatin state of ETS motif-containing promoters was preferentially affected upon loss of CHD8. Among the many ETS transcription factors, we found ELK1 to be the best correlated with CHD8 expression in primary human fetal and adult cortical neurons and most highly expressed in our ES cell-derived neurons. Remarkably, ELK1 was necessary to recruit CHD8 specifically to ETS motif-containing sites. These findings imply the functional cooperativity between ELK1, a key downstream factor of the MAPK/ERK pathway, and CHD8 on chromatin involvement in human neurons. THEREFORE, the MAPK/ERK/ELK1 axis may also play a role in the pathogenesis caused by *CHD8* mutations ^5^.

## Main

To study the role of CHD8 in human neurons, we engineered the *CHD8* locus in pluripotent stem cells to produce heterozygous and homozygous conditional knockout (cKO) cells. The heterozygous cKO allele was constructed by surrounding exon 4 with two loxP sites (Fig. 1a, Extended Data Fig. 1a). Deletion of exon four is predicted to produce a frameshift and early termination mutation. We identified two correctly targeted embryonic stem (ES) and one correctly targeted induced pluripotent (iPS) cell line (Extended Data Fig. 1b-c). To generate a homozygous cKO of *CHD8*, we used CRISPR/Cas9 to introduce an indel mutation in the nontargeted wild-type allele of heterozygous cKO cells (Fig. 1b, Extended Data Fig. 1h, i). This effort resulted in two homozygous cKO ES and one iPS cell line (Extended Data Fig. 1j). To conditionally deplete CHD8 protein, we infected heterozygous and homozygous cells with lentiviral vectors encoding Cre recombinase or ΔCre- an inactive recombinase, and we confirmed successful protein depletion by Western blotting and immunofluorescence (Fig. 1d, Extended Data Fig. 1g,k). Importantly, depletion of CHD8 in differentiated neurons did not affect overall cell viability (Extended Data Fig. 1k). Therefore, we could conduct an in-depth characterization of the cell biological consequences of loss of CHD8 in human neurons.

**Fig. 1:**
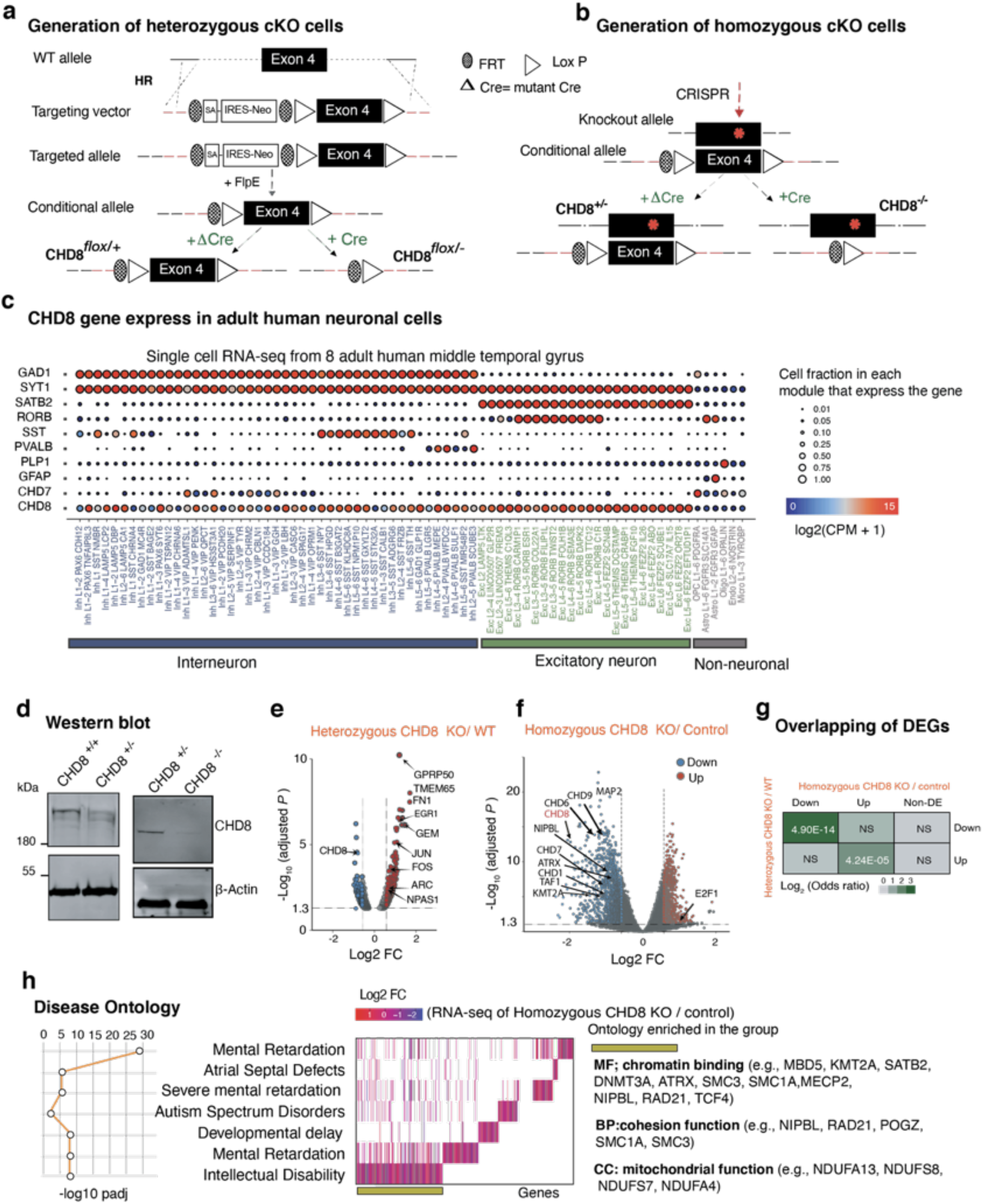
Widespread transcriptional changes upon loss of CHD8 in neurons. **a,** Strategy to generate a heterozygous conditional knockout (cKO) allele of *CHD8* in pluripotent stem cells. The endogenous Exon 4 was flanked with LoxP sites by AAV-mediated homologous recombination. Following correct targeting, the selection cassette was removed by transient transfection with FlpE recombinase to generate the final conditional allele. Infection with Cre recombinase leads to deletion of the floxed allele and generates CHD8 KO cells. Infection with ΔCre (an inactive form of Cre) is used throughout the study for the control condition. **b,** Generation of homozygous *CHD8* cKO cells by introducing a CRISPR transfection-mediated indel mutation into the non-conditional *CHD8* allele, which led to a frameshift mutation. Infection of a correctly targeted line with Cre recombinase generates homozygous CHD8 null cells, whereas control infection with ΔCre leaves the engineered mutations unchanged. **c,** Single-cell RNA sequencing data showing expression of CHD8 and cell-type-specific marker genes (CPM+1) in human cortical neurons (image credit: Allen Institute)^6^. **d,** Western blot from conditional heterozygous and homozygous KO neurons show a decrease and a near-complete depletion of the protein in each system. **e,** Volcano plot for RNA-seq fold change in heterozygous CHD8 KO vs. control neurons. **f**, Volcano plot for RNA-seq fold change in homozygous CHD8 KO vs. control neurons. **g,** Analysis for overlapping DEGs between the heterozygous and homozygous knockout RNA-seq experiment shows that the odds ratio of downregulated genes is significantly higher compared to overlapping upregulated genes. **h,** Disease Ontology (DO) and enrichment analysis for DEGs from homozygous RNA-seq experiment show enrichment of genes for human diseases (left). Additionally, it shows the association of molecular and biological ontology within the group of genes involved in the pathology of intellectual disability (right)^23^.

Analysis of single-cell RNA sequencing data from the middle temporal gyrus of human cortex ^6^ revealed that *CHD8* is predominantly expressed in excitatory neurons among other cell types investigated (Fig. 1c). Therefore, we characterized conditional *CHD8*-mutant cells in excitatory neurons using our previously published Ngn2 differentiation method ^7^. The basic electrophysiological properties of CHD8-mutant neurons were not obviously altered. Intrinsic membrane properties of neurons were unchanged in both heterozygous and homozygous *CHD8*-mutant cells (Extended Data Fig. 1l, p). Active membrane properties induced by stepwise current injection were similar between the heterozygous and homozygous mutants and the respective control cells (Extended Data Fig. 1m, q). Additionally, the frequency and amplitude of spontaneous miniature EPSCs in CHD8 heterozygous cKO cells were not statistically different from WT neurons (Extended Data Fig. 1o). Similarly, we found that evoked excitatory postsynaptic currents (EPSCs) were unchanged in heterozygous and homozygous mutant cells compared to respective control cells (Extended Data Fig. 1n, r). Thus, loss of *CHD8* did not grossly affect the intrinsic physiological and basic functional synaptic properties of human neurons using standard electrophysiology.

Given CHD8’s proposed role as chromatin regulator, we next evaluated the transcriptional effects following CHD8 depletion ^8^. Quantification of gene expression by RNA-sequencing showed that heterozygous *CHD8* mutant cells exhibited only subtle changes compared to WT cells, as described before (Fig. 1e) ^9,10^. Gene expression changes were much more pronounced in homozygous *CHD8*-mutant neurons (Fig. 1f). Overall, downregulation of genes was more pronounced than upregulation in the homozygous KO cells, and more DEGs (differentially expressed genes) were downregulated than upregulated (Fig. 1f). The overlap between heterozygous and homozygous KO cells was also higher among the downregulated genes (Fig. 1g). The enrichment analysis for molecular, cellular, biological, and disease pathways showed a significant overrepresentation of pathways involved in molecular regulations of chromatin, transcription, and pathways related to ERK/MAPK signaling and cell adhesion molecules (Extended Data Fig. 1s). Accordingly, disease pathway enrichment and disease-associated Gene Set Enrichment Analysis (GSEA) analysis showed an enrichment of gene signature for neurodevelopmental diseases and ASD (Fig. 1h, Extended Data Fig. 1t,u)^11^.

To map the chromatin targeting of CHD8 in human neurons, we generated a human embryonic stem (ES) cell line in which we tagged the endogenous CHD8 gene with a C-terminal FLAG-HA-tag (Fig. 2a, Extended Data Fig. 2a-d). Western blotting showed that the tagged protein had the expected size of CHD8 (Fig. 2b). We performed chromatin immunoprecipitation followed by sequencing (ChIP-seq) upon differentiation of ES cells into neurons using antibodies for both HA and the N-terminus of the CHD8 protein. There was a good correlation between the ChIP-seq with the two antibodies (Pearson *r^2^* =0.80, Extended Data Fig. 2e,f). Conversely, the correlation of CHD8 binding with a published ChIP-seq dataset from neural progenitor cells was low ^10^ (Pearson *r^2^* =0.25) (Extended Data Fig. 1f). These observations suggest genomic binding of CHD8 is strongly context dependent.

**Fig. 2:**
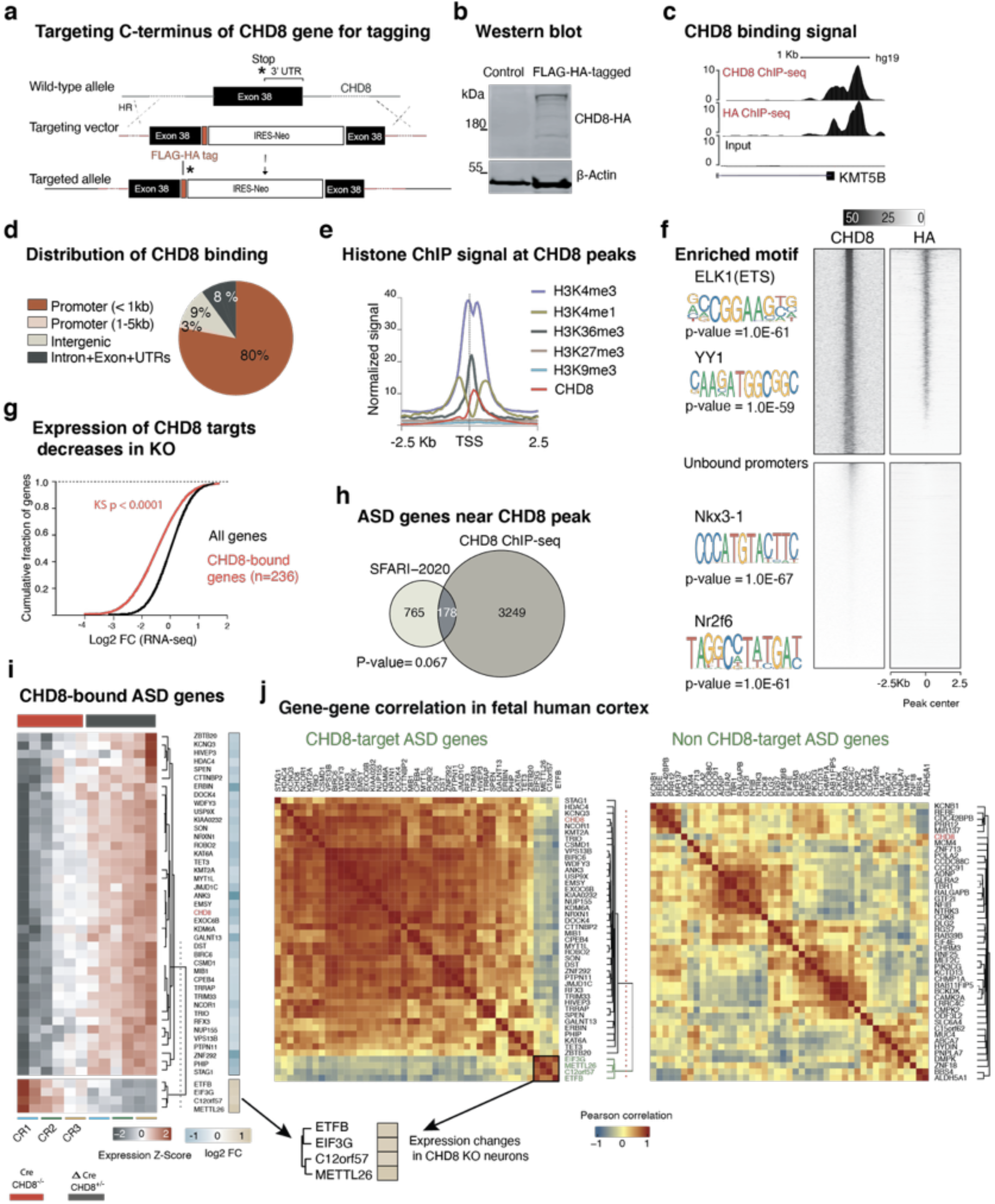
CHD8 localizes to promoters enriched with ELK1 motifs. **a,** Targeting strategy to insert a C-terminal FLAG-HA tag at the endogenous *CHD8* locus (see Extended Data Fig. 2a-d). **b,** Western blot analysis of tagged and non-tagged (control) ES cells using an anti-HA antibody. **c**, An example of a CHD8 peak at the promoter of the *KMT5B* locus. **d,** Pie chart shows the distribution of CHD8 ChIP-seq peaks across the genomic regions in ES cell-derived neurons. **e**, In neurons CHD8 binding sites enriched with active histone modifications (analysis of ENCODE data for H9 cell-derived neurons) ^13^. **f**, Top heatmaps are CHD8 binding at overlapping peaks between two separate pull-downs of HA and CHD8 antibodies (n=3696 overlapping peaks). Using HOMER and MEME, we found ETS and YY1 motifs enriched at CHD8-bound sites. The bottom panels’ heatmaps are signals from the same samples that stratified on CHD8-unbound promoters (n=4000). The greyscale shows normalized coverage for all groups ^24,25^. **g**, Expression of genes with CHD8 peak at the +/- 5Kb of the promoters showed a marked decrease compared to control. **h**, Overlapping of ASD genes from SFARI-2020 list and genes with CHD8 binding at promoters^11,26^. Significance of the overlapping calculated with the hypergeometric test. **i,** Gene expression of CHD8-bound ASD genes with significant expression change in RNA-seq experiment. **j,** correlation analysis of CHD8-target ASD genes (same gene set we discovered in “i”) within the human fetal cortex (12-37 CPW) is plotted. The clusters of similarly expressed genes led to the separation of the same gene modules shown in “i”. A separate module of non-CHD8 target autism gene was randomly selected for correlation analysis of the control group.

Classification of CHD8 binding revealed a distinct enrichment around the proximal promoters and little binding at distal regulatory sites (Fig. 2d, Extended Data Fig. 2g)^12^. Further characterization employing ENCODE repository of histone ChIP-seq data from H9-derived and human primary neurons revealed that the sites of CHD8 binding promoters (n=3,600) overlap with active histone marks (Fig. 2e, Extended Data Fig. 2i,j). The ontology of the associated genes with those promoters is related to regulating chromatin, transcription, and translation (Extended Data Fig. 2h) ^13^.

Motif enrichment analysis showed that CHD8 binding sequences in human neurons are enriched for the ETS and YY1 motifs, and the odds ratio of the enrichment of ETS motifs was significantly higher in the strong binding sites of CHD8 compared to weaker binding sites (Fig. 2f, Extended Data Fig. 2k)^14^. Non-CHD8 occupied promoters lacked such enrichment (Extended Data Fig. 2l, m).

We then investigated the functional consequences on the local chromatin at CHD8 target sites in response to CHD8 depletion. Changes activating or repressive binding of CHD8 by analyzing transcriptional changes of associated genes with CHD8-bound promoters in response to CHD8 depletion. RNA-sequencing between control and CHD8 KO cells in the cumulative distribution of CHD8-bound and unbound genes showed that loss of CHD8 leads to downregulation of its target genes, suggesting CHD8 primarily acts as a transcriptional activator of its direct target genes (Fig. 2g). Indeed, a comparison of odds ratios for CHD8 binding at promoters of up-regulated and down-regulated genes in heterozygous and homozygous mutant cells using RNA-seq validated the finding. Conversely, we also found that CHD8 binding is stronger at promoters of down-regulated genes than of up-regulated genes in both conditions (Extended Data Fig. 2n).

Among the CHD8 target genes that were predominantly down-regulated were many Autism risk genes defined by the Simons Foundation Autism Research Initiative (Fig. 2h, i, Extended Data Fig. 2o). Co-expression analysis of this set of genes in single-cell RNA-seq data of the human fetal brain revealed a striking separation of the genes up-and down-regulated following CHD8 depletion suggesting a functional relationship within these two groups of genes (Fig. 2j)^6^.

To characterize the presumed function of CHD8 as a chromatin remodeler in human neurons, we then performed Assay of Transposase Accessible Chromatin (ATAC)-seq to assess high-resolution chromatin accessibility of chromatin ^15,16^. Differential accessibility analysis revealed the vast majority represent a loss of accessibility in CHD8-mutant cells: 1481 peaks lost accessibility, but only 106 peaks gained accessibility in KO neurons (Fig. 3a, b, see also the PCA analysis in Extended Data Fig. 3a). The ontology enrichment analysis for genes with a significantly changed ATAC-peak signal at their promoter vicinity (+/- 5 Kb) produced terms about regulation of transcription and RNA binding pathways (Fig. 3c). Motif enrichment showed there is an enrichment of CAAT (an RNA polymerase II binding sequence) and the GGAA (the ETS factor motif) at sites that lost accessibility in CHD8 knockout neurons (Fig. 3d). The CAAT-box enrichment is commonly found at core promoters ^17^. However, the enrichment of an ETS motif was intriguing since we had already found it to be enriched at CHD8 targets. We therefore next interrogated CHD8 binding at the dynamically regulated ATAC-seq sites. We found that CHD8 binding strongly enriched at differentially accessible promoters in particular promoters closing without CHD8 (Fig. 3e, f, Extended Data Fig. 3b). Overall, across the genome, CHD8 binding was enriched at all sites that showed differential chromatin accessibility in KO cells, compared to unchanged ATAC-seq sites (Fig. 3g). These results suggest that CHD8 regulates its chromatin targets through direct genomic interaction.

**Fig. 3:**
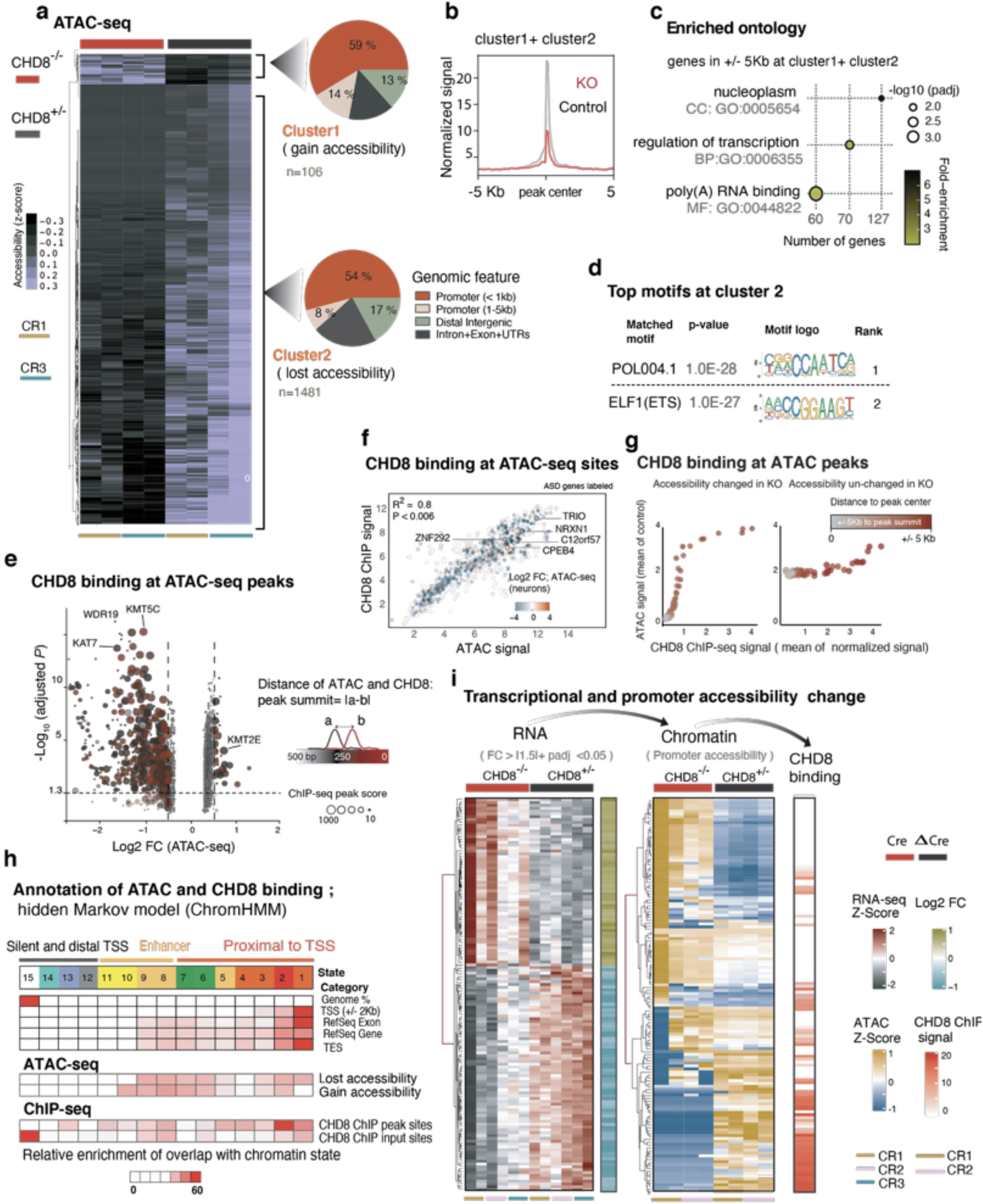
CHD8 activates chromatin at its targets in neurons. **a**, Heatmaps of normalized ATAC-seq signal in cKO homozygous and control neurons from a targeted ES cell line (CR1) and a targeted iPS cell line (CR3) and two technical replicates within each line. Cluster1 and Cluster 2 are separated based on unsupervised clustering analysis, which shows sites that gain or lost accessibility regions in KO and plotted the corresponding genomic annotation at each cluster. **b**, Normalized ATAC-seq signal from aggregates of Cluster1 and Cluster2 shows KO chromatin predominantly lost accessibility. **c**, Ontology analysis of the genes with differentially regulated ATAC-seq peaks in KO at the promoters’ vicinity (+/- 5Kb TSS in Cluster1 +Cluster2). **d**, Motif enrichment analysis in Cluster 2. **e**, Volcano plot depicting each ATAC-seq peak as one dot. The color indicates the distance of the ATAC-seq peak summit from the CHD8 peak summit at the immediate vicinity, and the size reflects the CHD8 ChIP-seq peak score. **f**, Linear regression of DE-seq2 normalized CHD8 binding signal and DE-seq2 normalized ATAC-seq signal at the promoters of genes with change accessibility, and example ASD genes are labeled. The color indicates log2 change of the accessibility in CHD8 KO. **g**, Analysis of CHD8 binding and ATAC-seq signal on sites with differential accessibility in *CHD8* KO neurons and sites with no change. The average ATAC-seq signal is calculated from control samples. **h,** Analysis of chromatin modification enrichment with ChromHMM and the annotation of the genomic feature with transition probability for 15 state model uncovers the distribution and the relative enrichment of CHD8 binding and the ATAC-seq sites across all chromatin states in neurons. **i,** RNA-seq and ATAC-seq signal for **136** genes with a significant change in gene expression and ATAC-seq signal in KO. CHD8 binding signal sorted with the same order of the sites of the promoters at the respective heatmap.

Next, we were investigating the relationship between CHD8-dependent chromatin accessibility, differential gene expression and CHD8 chromatin binding. Averaging fold changes of all genes proximal to opening and closing ATAC-seq peak showed changes in ATAC-seq signal in KO is at the same direction as the changes of the RNA expression at the corresponding gene (Extended Data Fig. 3c). This observation was also true for high-confidence autism genes as defined by SFARI (Extended Data Fig. 3d). These data indicated that chromatin changes in CHD8 KO cells are in fact relevant and affect gene expression in neurons. We then applied a multivariate hidden Markov model (HMM) to annotate the genome-wide chromatin state of CHD8 targets using publicly available datasets for chromatin modifications of human H9-derived neurons ^18,19^. First, we validated that our model accurately described the expected chromatin state at a group of well-annotated promoters (n=500) (Extended Data Fig. 3e). Next, we analyzed the enrichment of CHD8 targets, including the sites of CHD8 binding and the ATAC-seq peaks at annotated genome. Enrichment analysis revealed CHD8 regulates chromatin accessibility at regions of the genome with active chromatin state with no preference to a distinct classification or mapping to a particular genomic annotation (e.g., promoters or enhancer). In contrast, CHD8 binding displayed a strong preference for proximal promoters (Fig. 3h). To validate our annotation analysis, we analyzed CHD8 binding at the promoters of a distinct group of 136 differentially expressed genes that also in ATAC-seq KO their promoter accessibility changed. We observed CHD8 binding strongly enriched at closing promoters in downregulating genes (Fig. 3i). These results reveal CHD8 directly activates chromatin and RNA expression at its overlapping targets.

To further explore the possible role of ETS motifs guiding specificity of CHD8 interaction and effects on chromatin accessibility, we analyzed regulatory DNA sequence and motif elements in ATAC-seq target regions with or without ETS motifs. We interrogated the chromatin dynamics at the selected sites and observed a much more pronounced loss of ATAC-seq signal at the ETS motif containing sites compared to ETS motif-free regions (Fig. 4a). In combination with ETS motif enrichments in CHD8 binding sites and CHD8-dependent ATAC-Seq sites, these results suggested the possibility of a functional interaction of an ETS factor and CHD8. To further characterize ETS motif-dependent CHD8 activity, we implemented cross-correlation analysis of the ATAC-seq signal to infer nucleosome density at transcriptional start sites (TSS) with ETS motifs but not at TSS lacking ETS motifs (Fig. 4b, c)^16^. We found the altered density of the +1 nucleosome intriguing, prompting us to investigate the symmetry of CHD8 binding at promoters with and without ETS motifs. Indeed, the average CHD8 ChIP-seq signal (100 bp binned) upstream of promoters of actively transcribed genes in neurons was stronger than downstream of promoters, again specifically at ETS motif-containing sites but not others (Fig. 4d).

**Fig. 4:**
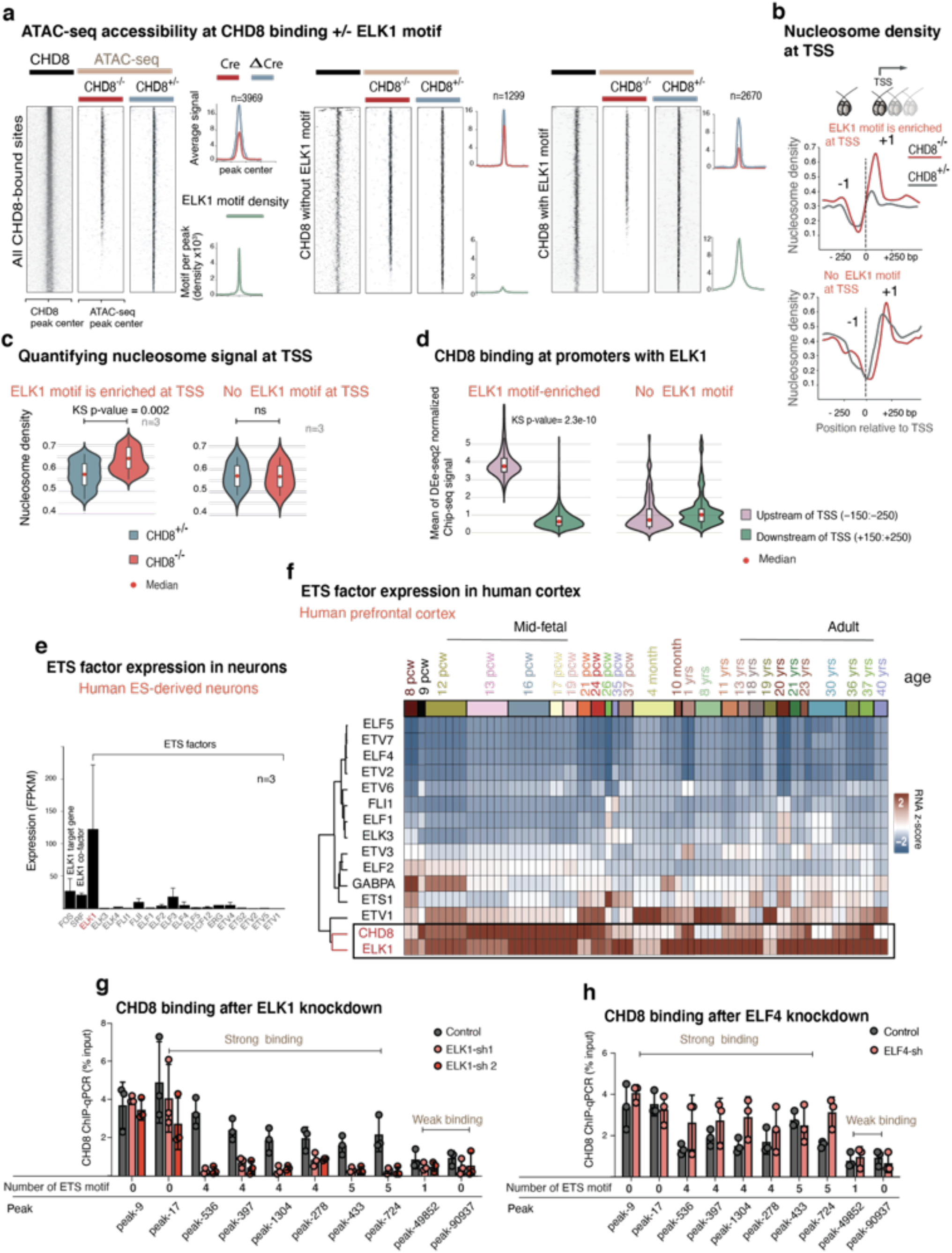
Demonstration of functional cooperativity between CHD8 and ELK1 in targeting and regulating chromatin. **a**, Normalized CHD8 signal and ATAC-seq signal in *CHD8*-KO and the control samples plotted, and ELK1 motif density plotted as a green enrichment plot. Genome-wide ChIP-seq and ATAC-seq signal divided into three groups: sites with exclusive enrichment for ELK1 motif, sites without ELK1 motif, and the control sites, which are the randomly shuffled peak sets from control samples. For each group, we compared the ATAC signal from KO to the control sample. **b,** Shows cross-correlation of ATAC-seq signal with NucleoATAC to measure nucleosome density. The enrichment plot is a subset of calculated nucleosome density signal taken from TSS with +/- ELK1 (ETS) motif (motif occurrence >1).**c**, Violin plots compare normalized and averaged nucleosome density signal at 100bp region around the position +1 of TSS with the presence or absence of ELK1 motif. Statistical analysis of distribution comparison is calculated with the Kolmogorov Smirnov test. **d**, CHD8 binding signal taken from 100 bp upstream and downstream of transcription start sites (TSS), with the presence or absence of ELK1 motif. **e,** Average expression of ETS factors shows ELK1 is the only highly expressed ETS gene in differentiated human neurons (average FPKM values taken from wild type neurons). **f**, Gene expression of *CHD8* and ETS factor analysis in human developing human cortex and unsupervised clustering of expression levels (Alan brain data)^6^. **g**, ChIP-qPCR analysis for CHD8 binding after ELK1 is KD with two different hairpin RNAs (shRNAs). The control condition is the empty vector. The number of ETS motif at each peak site indicated beneath each peak. There was no change in CHD8 binding at sites without the ETS motif. See also Extended Data Fig. 4d for validation of CHD8 binding on the peaks in CHD8-KO neurons. **h**, Knockdown of ELF4 does not affect CHD8 binding on either of ETS or YY1 motif sites, suggesting ELF4 does not influence chromatin binding of CHD8.

Which ETS factor may functionally interact with CHD8 in human neurons? First, we turned to our gene expression data from wild-type neurons. We found that among all *ETS* factors *ELK1* is most highly expressed in human neurons (Fig. 4e) ^6,20^. Next, we analyzed the expression of *ETS* factors in human prefrontal cortex. Clustering analysis of single cell RNA-seq data from the human prefrontal cortex revealed a correlative gene expression pattern between *ELK1* and *CHD8* but no other *ETS* factors (Fig. 4f). These results point to the ELK1 transcription factor as potential functional partner of CHD8 and ELK1.

For directly assess the functional cooperativity between ELK1 and CHD8 in targeting chromatin, we constructed lentiviral vectors with short hairpin RNA (shRNA) targeting *ELK1* and *ELF4* as control. We measured the binding of CHD8 at a series of CHD8 binding, ETS motif-enriched peak regions (see the experimental diagram at the Extended Data Fig. 4a). Quantitative qPCR and western blotting confirmed a robust decrease in mRNA and the protein after infecting the neurons with two hairpins against *ELK1* and one hairpin against *ELF4* (Extended Data Fig. 4b). We also validated the specificity of our selected ChIP-seq peaks for CHD8 binding in three independent pull-down experiments, which showed a complete absence of CHD8 binding in KO neurons (Extended Data Fig.4c, d). After these validation experiments, we then asked whether ELK1 would be required for proper CHD8 targeting and chromatin binding. To that end, we performed ChIP on selected sites based on strength of CHD8 binding and number of ETS motifs. The results were striking: Among all strong CHD8 binding sites interrogated that also contained ETS motifs, ELK1 knockdown reduced CHD8 binding (Fig. 4g). In contrast, CHD8 peaks that did not contain ETS motifs were unaffected. The knockdown of the other ETS factor *ELF4* did not change CHD8 binding at the same peak sites (Fig. 4h). Thus, ELK1 is the critical ETS factor necessary for the proper chromatin targeting of CHD8.

In summary, these results establish that CHD8 is responsible for maintaining an open chromatin configuration and overall transcriptional activation in human neurons. Our data also reveals functional cooperativity of *CHD8* and *ELK1* (the effector of MAPK/ERK) in chromatin regulation through a distinct model of directional activity oriented around the ELK1 motif. Additionally, we revealed CHD8 regulates a distinct group of autism genes positively correlated in expression patterns in developing human cortex, suggesting a conserved and developmentally regulated transcriptional connectivity between CHD8 and its targets. In light of these data, it is intriguing to speculate that MAPK/ERK/ELK1 may play a functional role in developing neuropsychiatric alterations caused by *CHD8* mutations. Modulation of specific aspects of this pathway, which is known to regulate activity-dependent gene expression and synaptic plasticity, may represent a foundation to explore a therapeutic opportunity for functional interference with pathology induced by *CHD8* mutations ^21,22^.

## Acknowledgments

We would like to thank Dr. Joanna Wysocka, Dr. Tomasz Swigut, and Dr. Daniel Jarosz for support and thoughtful discussions. This project was supported by an Howard Hughes Medical Institute Faculty Scholar Award and a New York Stem Cell Foundation Investigator Award (M.W.). T.C.S. and H.Y.C. are Investigators of the Howard Hughes Medical Institute.

## Extended Data Figure Legends

**Extended Data Fig. 1.**
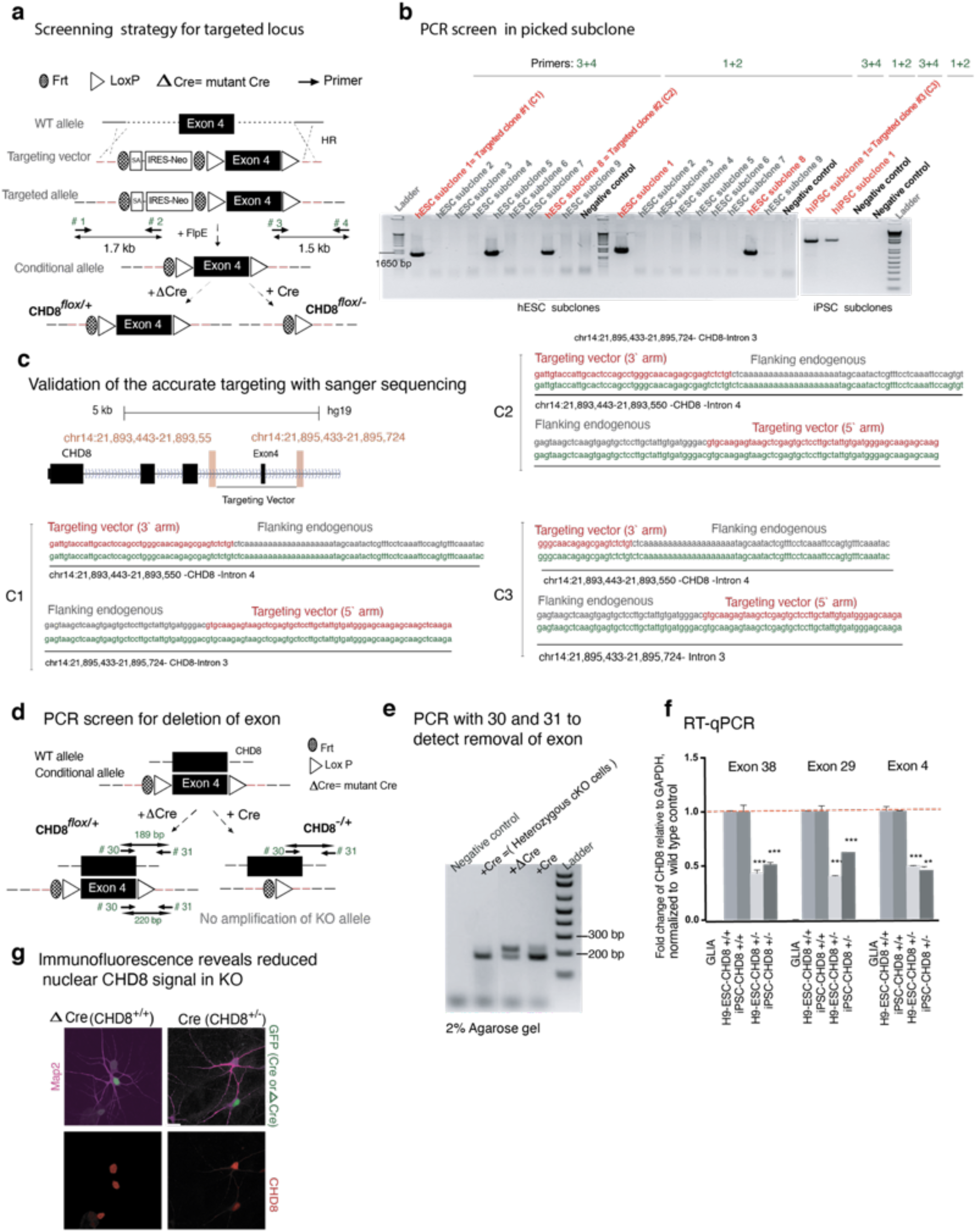
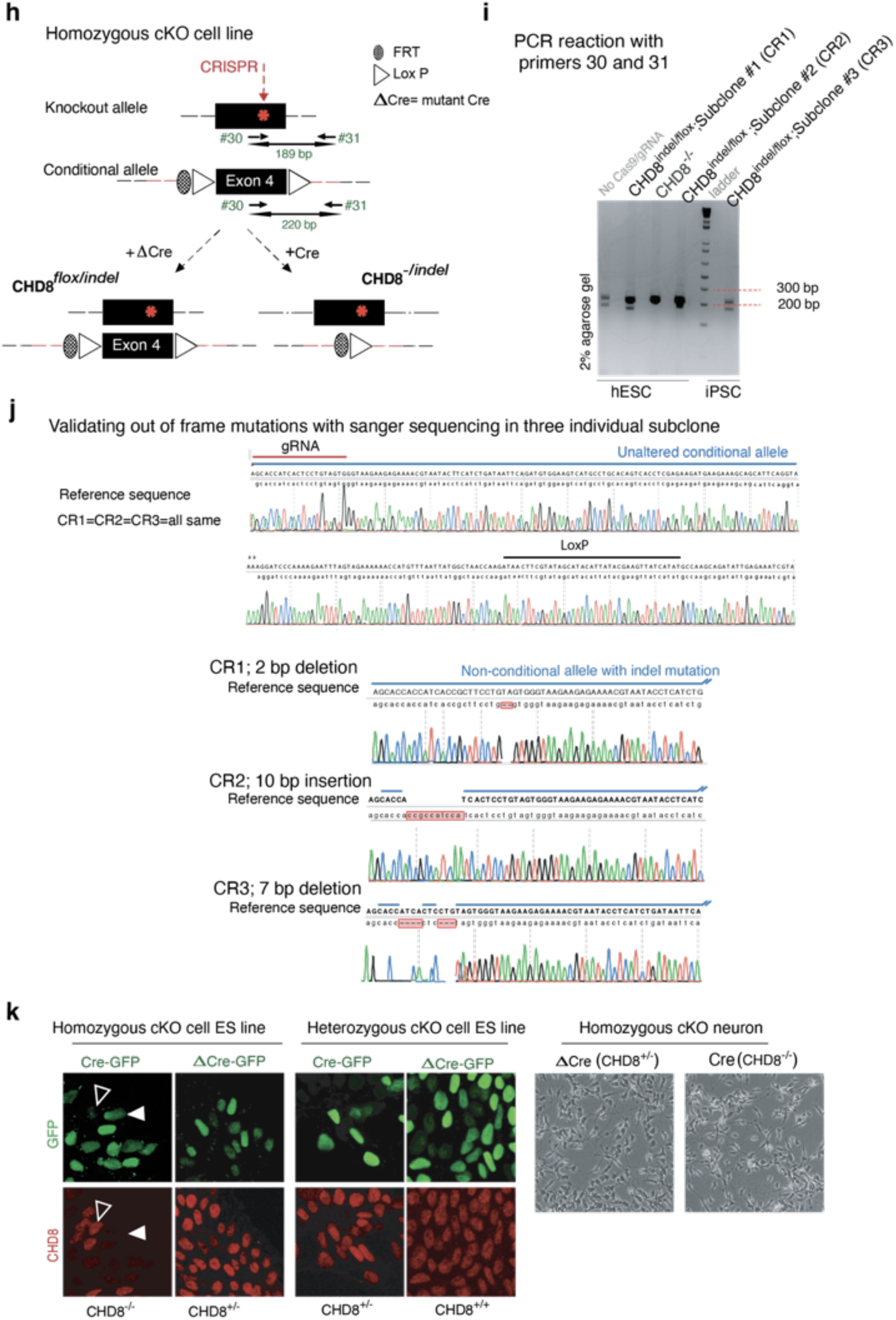
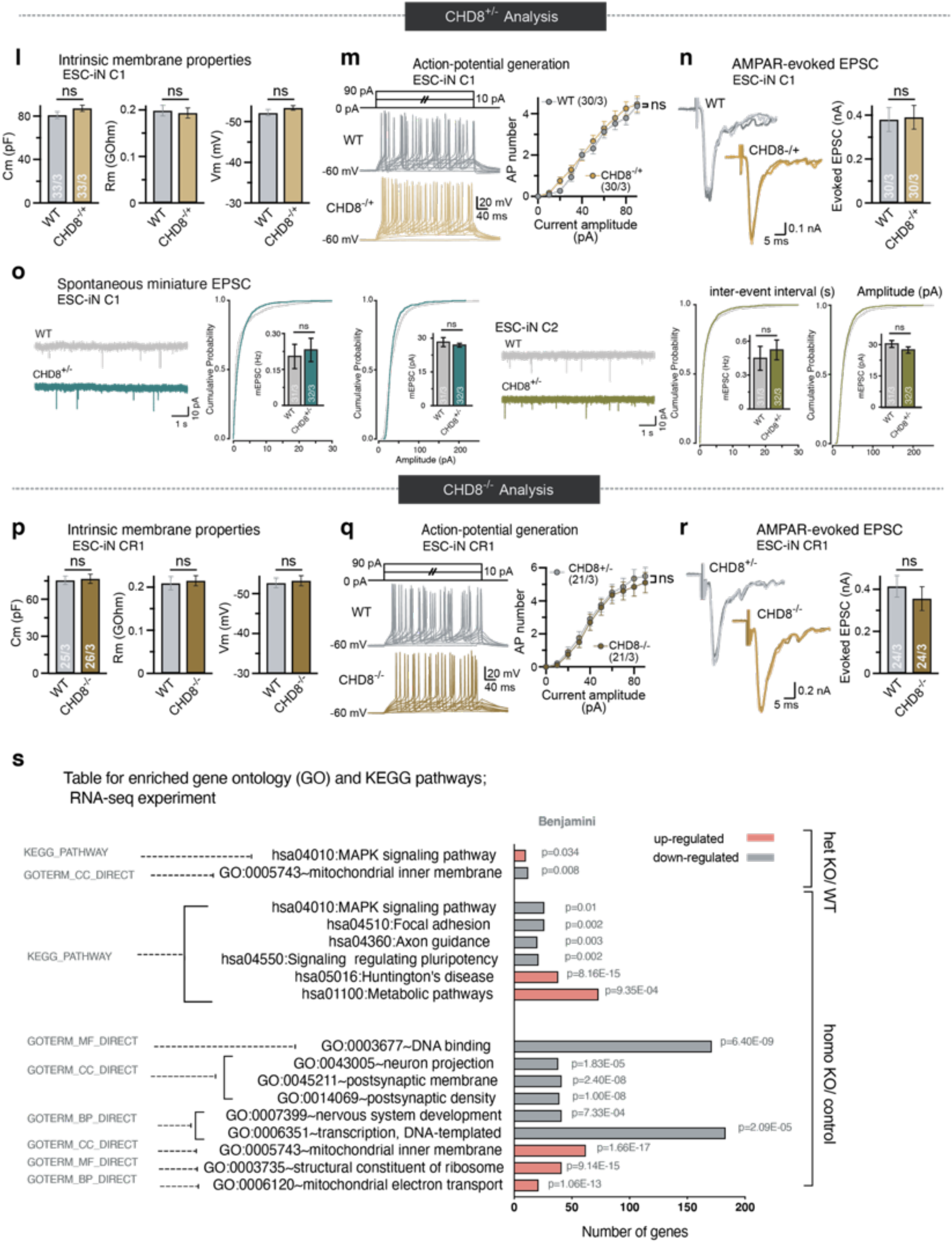
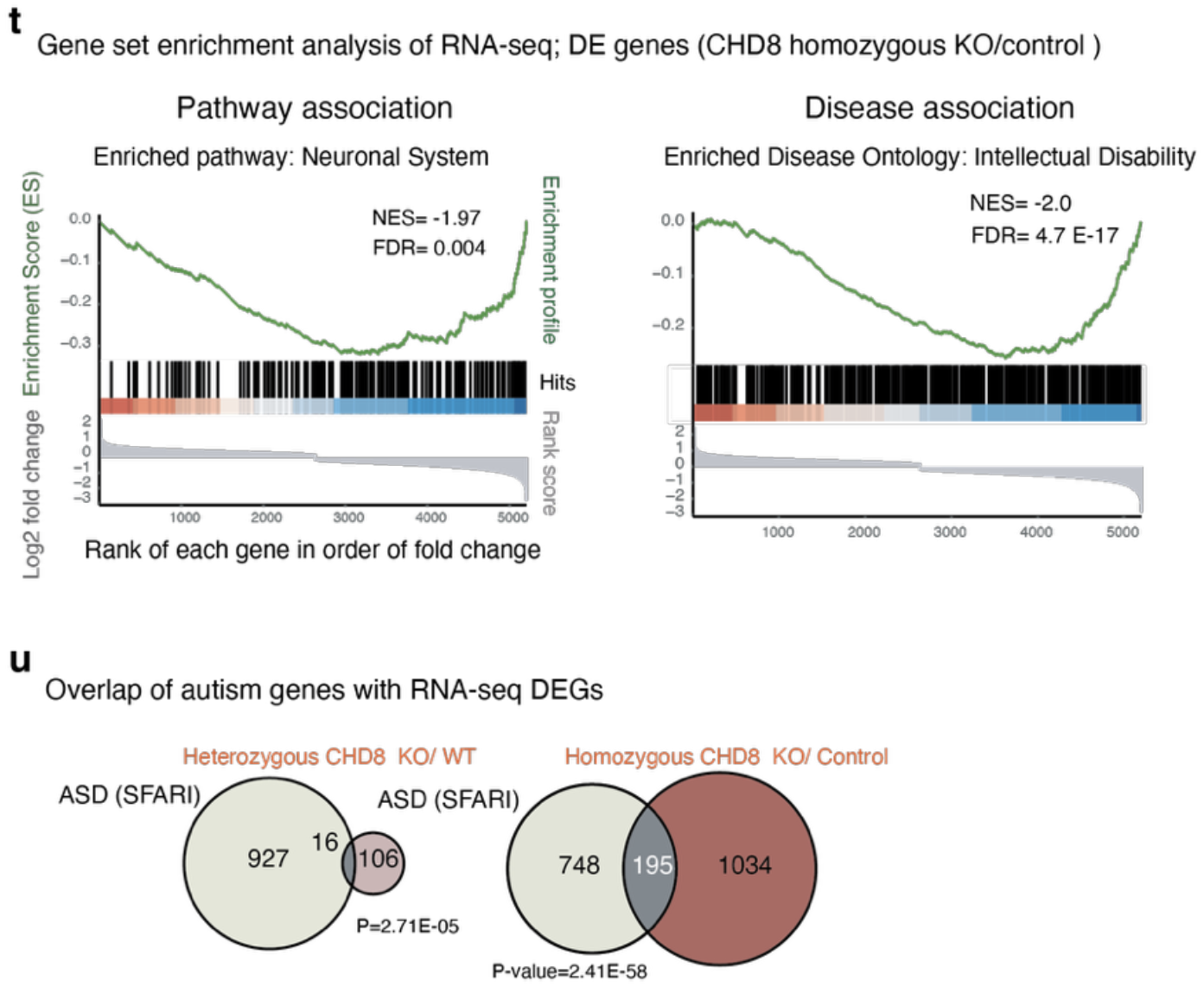
Targeting *CHD8* gene in human pluripotent stem cells. **a,** Schematics of Targeting strategy for generating a conditional KO allele of the *CHD8* gene by flanking exon 4 with LoxP sites. Deletion of exon four is predicted to create a frameshift mutation with early truncation. **b**, Screening PCR using external primers designed for outside the homology arm towards inside the targeting vector identified two subclones from the hESC line (C1 & C2) and one subclone from the iPSC line (C3) that were positive for the insertion of the targeting vector. **c**, Sanger sequencing is spanning the targeting arms’ transition into endogenous sequences, demonstrating correct targeting of the construct into the *CHD8* locus (clones C1, C2, and C3). **d**,**e**, Excision of exon four after infection with LV-Cre and screening with the primers around the loxP sites (primer 30 and primer 31) resulted in a single band from heterozygous KO compared to two bands in WT cells as expected. **f,** Quantitative reverse transcription PCR (RT-qPCR) using the probes for three exons of CHD8 gene shows the levels of mRNA decreases in heterozygous KO neurons. **g**, Immunofluorescence analysis of heterozygous KO and WT neurons for Map2 and CHD8. The nuclear staining signal intensity significantly decreases in heterozygous mutant neurons. **h,** Introduction of an indel mutation by CRISPR-CAS9 to non-conditional exon 4 of *CHD8* gene, to generate a conditional homozygous knock out cells. **i**, Validation of the genotype by PCR around the loxP sequence (spanning the gRNA targeting region) and amplification of two bands; one allele is 32 bp smaller than the other allele. Therefore, the top band corresponds to the floxed allele, and the bottom band is the non-conditional allele, a candidate for carrying an indel mutation. Each band is cut and gel-purified, TOPO cloned, and sequenced using M13 forward and M13 reverse primers (CR1, CR2; both hESC and CR3, is an iPSC subclone, confirmed to carry an indel mutation in non-floxed allele). **j**, Sanger sequencing of floxed and non-floxed alleles identified three subclones that carry frameshift indel mutations in the non-conditional allele with an un-altered floxed allele. Note that the conditional exon is shown only once as the representative sequence for all three subclones. **k**, Left shows immunofluorescent images of CHD8 staining in conditional heterozygous and homozygous *CHD8* KO embryonic stem cells three days after infection with Cre/Δ Cre to detect CHD8 reduction. Right depicts the bright-field images of homozygous *CHD8* KO and control neurons, 23 days after in vitro differentiation assay. **l,** *Electrophysiological* recordings of the intrinsic membrane properties in the C1 neurons, a heterozygous conditional KO line: capacitance (Cm), resting membrane potential (Vm), and input resistance (Rm). N=33 recorded cells in 3 batches (see numbers in bars). Student’s t-test. **m**, Active membrane properties of C1 neurons demonstrated by stepwise current injection protocol. The number of action potentials in response to current amplitude is plotted (right). **n**, Amplitude of evoked excitatory postsynaptic currents (EPSCs) in clone C1 showed no changes between the heterozygous KO and the WT neurons. **o**, Amplitudes and frequency of spontaneous miniature EPSC (mEPSCs) in the presence of 1μM tetrodotoxin showed no change between *CHD8* heterozygous KO and WT neurons in clone C1 and clone C2, N=31 or 32, respectively in 3 batches. Student’s t-test. **p**, Analysis of the conditional homozygous mutant cell line CR1 neurons. Shown are capacitance, input resistance, and resting membrane potential. **q**, Active membrane properties of CR1-derived neurons as in b. **r**, Recording of evoked excitatory postsynaptic currents (EPSCs) CR1 derived neurons shows no statistically significant difference between Cre and ΔCre (*CHD8*^-/-^ vs. *CHD8*^+/-^) neurons. N=24 cells in 3 batches, Student’s t-test. **s**, Results of gene ontology (GO) and KEGG pathway analysis for up and downregulated genes in heterozygous *CHD8* KO to WT and homozygous *CHD8* KO to control conditions. **t,** Gene set enrichment analysis of pathways and disease in *CHD8* homozygous KO to control conditions. **u,** Venn diagrams reveal overlapping CHD8-bound genes (ChIP-seq peak at +/- 5 Kb of the promoters).

**Extended Data Fig. 2.**
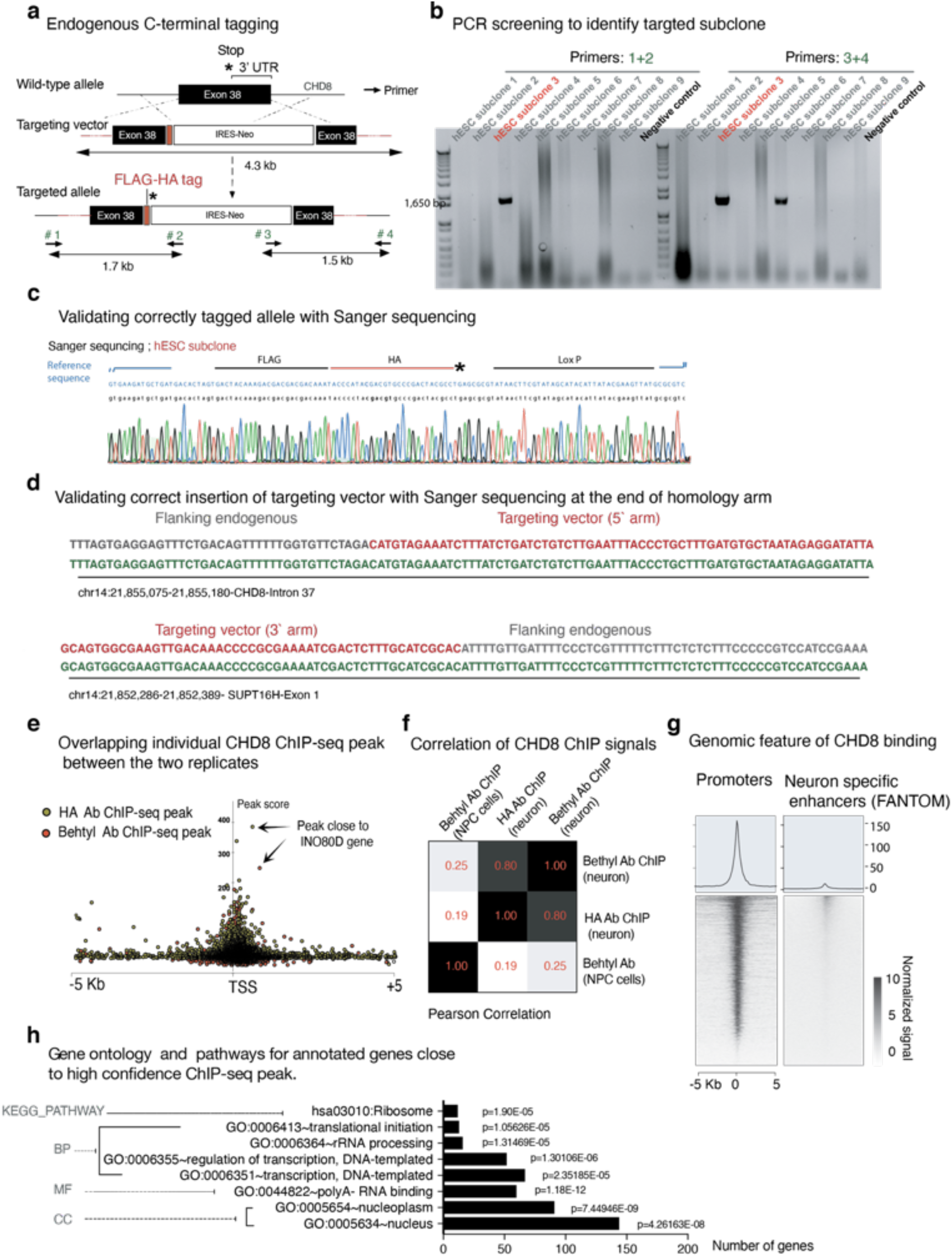
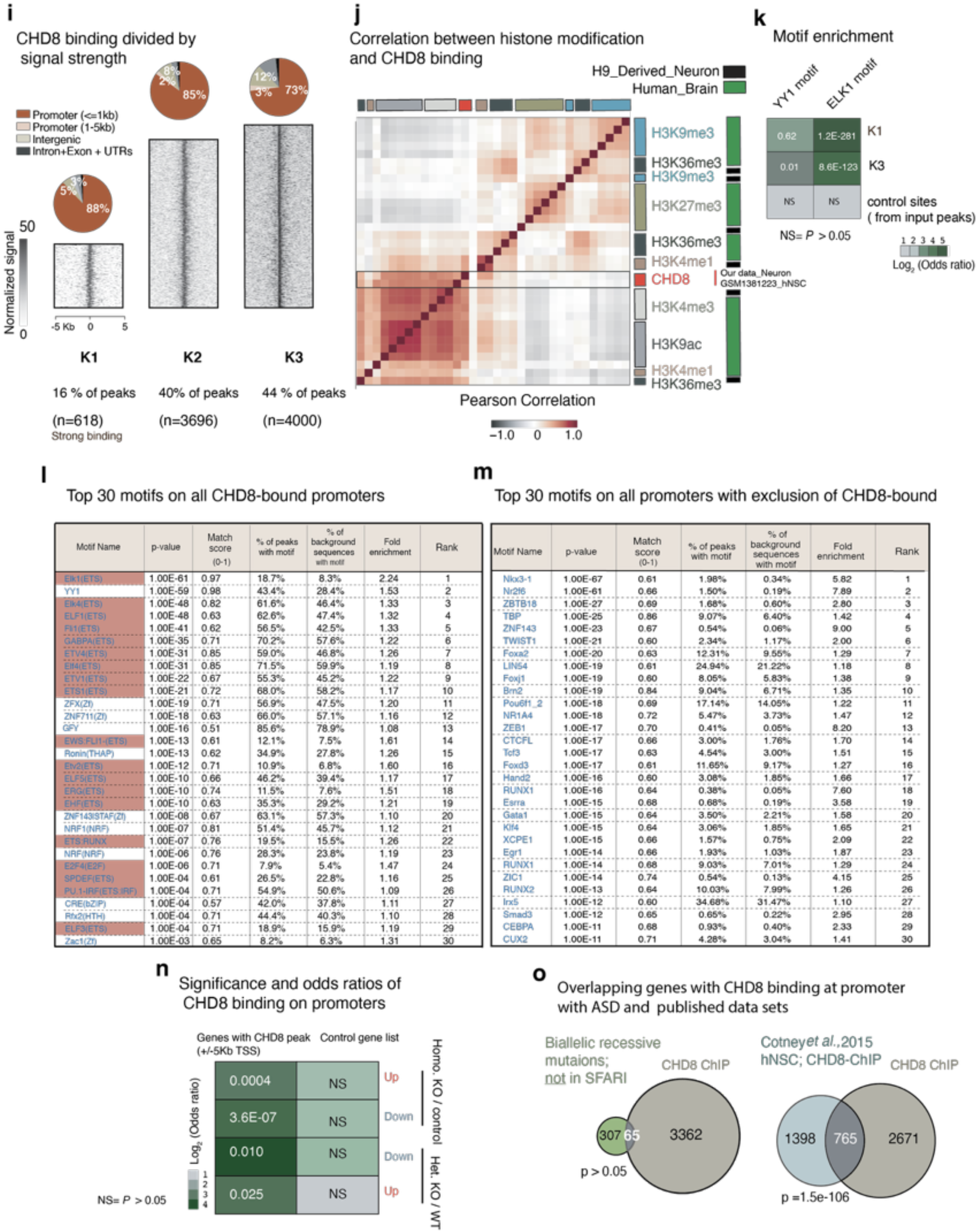
Targeting the C-terminus region of *CHD8* gene for endogenous protein tagging and ChIP-seq experiment. **a,** Schematic shows targeting strategy for in-frame insertion of FLAG-HA tag at the C-terminus region of CHD8 gene. Throughout the manuscript, we have used the HA tag for downstream experiments. **b,** Screening PCR of Neomycin resistant hESC colonies with external primers, i.e., one primer outside of the homology arms and one primer inside the targeting vector (primer #1 and #4). **c**, Sanger sequencing to detect a correct insertion of donor vector in the C-terminus of CHD8 gene within its frame and before the STOP (*) codon. **d**, Sanger sequencing to validate the correct transition of targeting vector into endogenous arms (black line after the blue lines on both sides). **e**, Individual CHD8 ChIP-seq peaks from two replicates that were pulled down with two different antibodies (anti-CHD8 and Rabbit anti HA). **f**, Pearson’s correlation between our two CHD8 ChIP-seq signals in neurons and a previously published CHD8 ChIP-seq dataset in NPC cells on selected, overlapping peak sites (GSE61492)^27^. Signals are compared on a set of selected, overlapping peak sites (10,000 peaks). **g,** Heatmap of CHD8 binding signal, at promoters of neural genes and enhancers. **h,** enrichment of pathway and the ontology of genes with CHD8 peak at their promoters. **i,** K-means clustering ChIP-seq signal into three groups recognize the most specific targets of CHD8 binding at promoters. Genomic annotation and functional classification are separately assigned for each group. **j,** Pearson’s correlation of histone modification from H9 derived human neurons (ENCODE’s histone ChIP-seq data) and our CHD8 ChIP-seq signal at CHD8 binding sites (K1+K2+K3). **k,** The odds ratio calculations of motif enrichment at strong CHD8 binding sites (K1), the weak binding sites (K3), and the control sites (randomly shuffled peaks from input). Sites (K1, K3) are taken from clustering analysis in Extended Data Figure 2i. **l,** Top 30 motifs that enriched at the prompters (+/- 5Kb) of 3000 CHD8-bound target genes. The ETS factor motifs are highlighted in red. **m,** Top 30 motifs that are enriched at 4000 control prompters of actively transcribed genes in neurons. **n,** The odds ratio, strength and significance of CHD8 binding on DEG promoters in heterozygous and homozygous CHD8 KO neurons (DE genes are obtained from RNA-seq experiment and odds ratio calculated based on overlapping gene list analysis) (GeneOverlap: http://shenlab-sinai.github.io/shenlab-sinai/). **o,** The overlapping autism-candidate genes that carry recessive gene disruption in the disease and a previously published data set of CHD8 binding in human NPCs with CHD8 target genes (our ChIP-seq experiment) ^28,29^.

**Extended Data Fig. 3.**
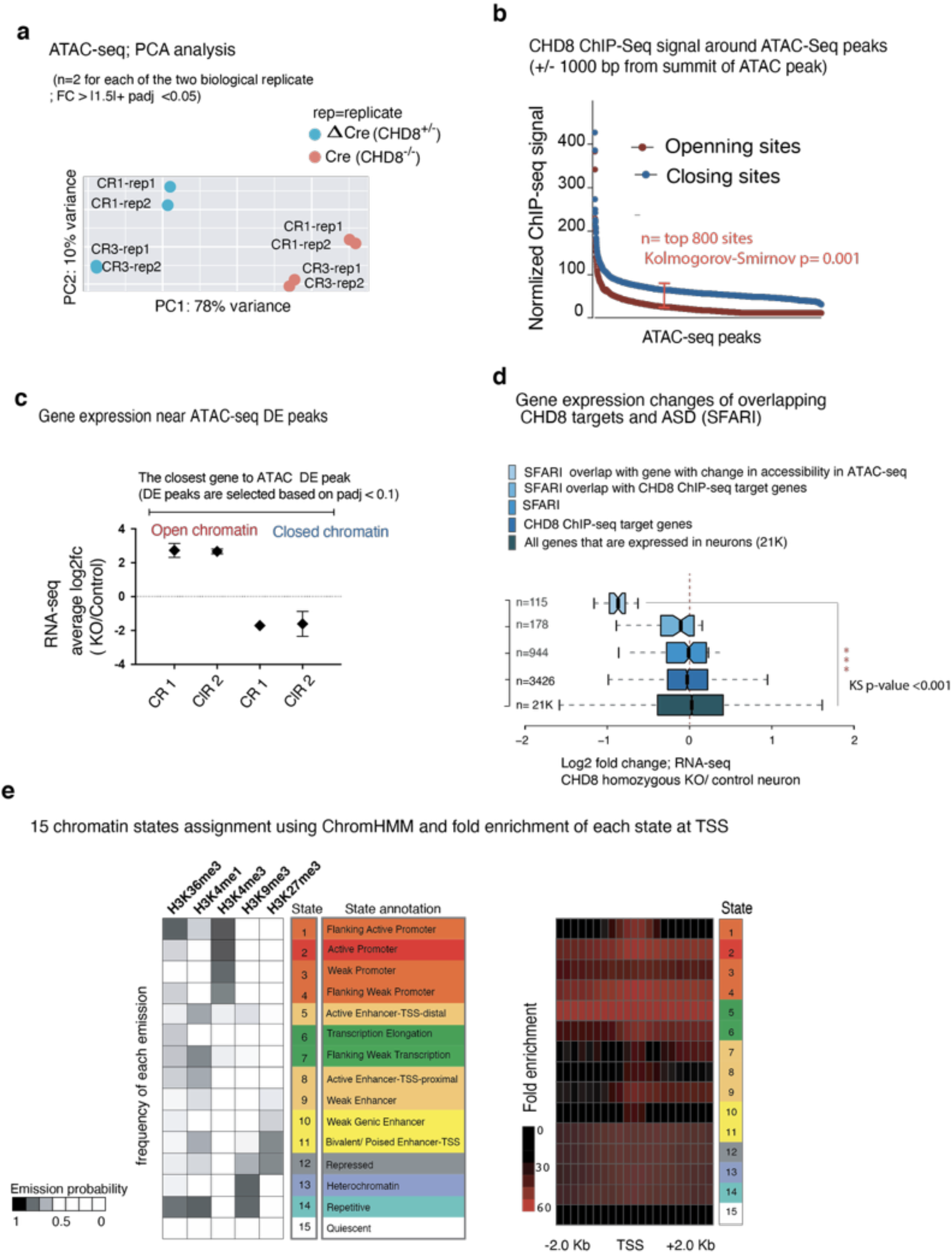
ATAC-seq experiment and the analysis of CHD8 target gene expression and the chromatin state of CHD8 target sites. **a,** Principal component analysis (PCA) for ATAC-seq signal from control (ΔCre) and homozygous KO (Cre) neurons and the separation of the samples based on the genotype in PC1. One embryonic stem cell line (CR1) and the One iPSC line (CR3), and two technical replicates for each line are used in the experiment. **b,** CHD8 binding on sites of chromatin that exhibit accessibility change in ATAC-seq experiment (CHD8 signal is taken from +/- 1000 bp of the summit of ATAC-seq’s DE peak). **c,** Gene expression of genes with open or close chromatin in CHD8-KO. **d,** Boxplots represent log2 scaled fold change of RNA expression of CHD8 target genes including overlapping with autism gene list and control gene list. **e,** Chromatin state analysis**: The right** heatmap represents an enrichment of promoter-associated state at annotated TSS regions. **The left plot** shows enrichment of histone signal (ENCODE data from H9 derived neurons) for each annotated state as the emission probability value.

**Extended Data Fig. 4.**
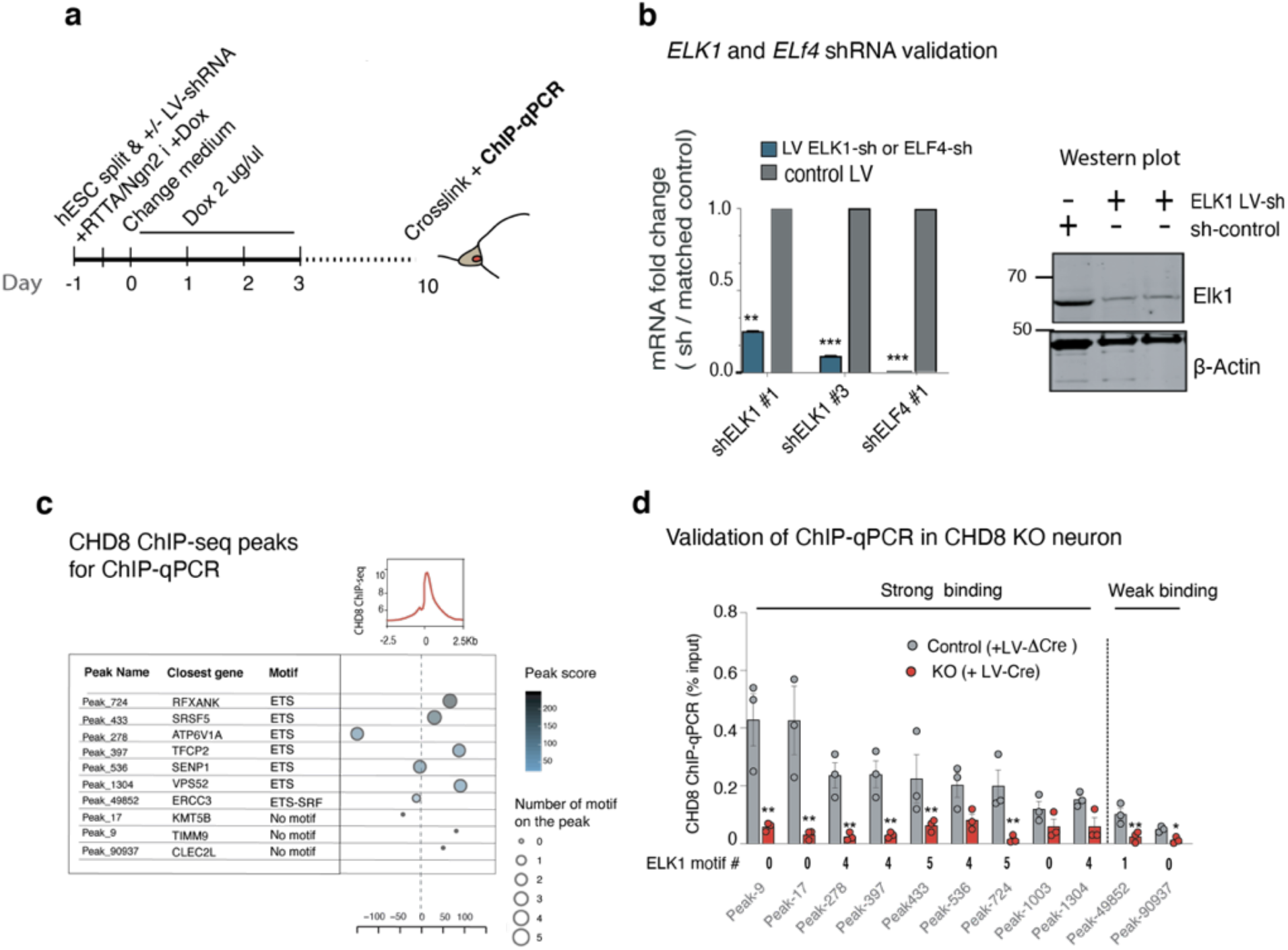
Analysis of CHD8 binding in *CHD8* KO neurons and selection of CHD8 peaks for interrogation of CHD8 binding in the absence of ETS factor. **a,** Schematics of shRNA gene knockdown and ChIP-qPCR experiment. **b,** Knockdown of *ELK1* and *ELF4* gene with a short hairpin (shRNA) shows a decrease in total mRNA (bar graph) and a significant reduction of ELK1 protein (western blot). **c,** Table of selected CHD8 ChIP-seq peaks, selected for ChIP-qPCR assessment; the number of motifs, peak score, and the location of the peak relative to TSS is shown. **d,** Validation of CHD8 binding on ChIP-seq peaks in CHD8 knockout neurons. Strong or weak peaks lost CHD8 binding in KO neurons, regardless of the motif at peak.

## METHODS

### Cell culture

*CHD8*-KO human ES cells were generated from the human embryonic stem cell (ESC) line H9 (passage 50, WA09 WiCell Research Institute, Inc.) and an iPSC line from a male individual. Only cells with normal karyotype were used to generate conditional knockout cells and downstream analysis. Pluripotent stem cells were maintained in mTeSR1 (STEMCELL Technologies), and small-molecule Thiazovivin (5μM) (STEMCELL Technologies) applied to the medium before single-cell passaging. The conversion of PSC to induced neurons is described below according to our previously published protocol ^7^.

### Lentivirus generation

Production of lentivirus was according to the previously described method ^30^.

### Production of Adeno-Associated Virus (AAV)

Recombinant adeno-associated virus (rAAV-DJ) was used to deliver the targeting vector to pluripotent stem cells. To produce rAAV we co-transfected three plasmids: 25 μg of pAAV ^31^, 25 μg of helper plasmid (pAd5) and 20 μg of capsid (AAV-DJ), into one T75 flask with 80% confluent HEK293T cells (ATCC) by calcium phosphate transfection method ^31,32^. Two days after transfection, cells were harvested by trypsin for 10 minutes and lysed by three rounds of freeze and thawing in dry ice and water bath (37 C). The rAAV virus was collected from the supernatant by spinning the whole lysate and removal of the pellet. The virus was aliquoted in small volumes to freeze in −80°C. Before usage for every 100 μl of supernatant, ten units of Benzonase endonuclease (EMD Chemical Inc, Merck 1.01695.002) added at (37°C) for 5 minutes to digest DNA from HEK cells; the capsid protects AAV DNA from digestion.

### Generation of human induced excitatory neurons (iN)

Human excitatory neurons differentiated from pluripotent stem cells by over-expression of lineage-specific transcription factor-Neurogenin 2 (Ngn2) as described before^7^. In summary, one day prior to conversion, we dissociated stem cells into single cells with Accutase (Innovative Cell Technologies) and seeded at ~ 40K cells into one 24 well plate pre-coated with Matrigel (BD Biosciences) in medium supplemented with Thiazovivin (5μM) (STEMCELL Technologies) and doxycycline (2 mg/ml, Clontech). After 6 hours, we infected the cells with lentivirus containing Ngn2, RTTA, and Cre recombinase or ΔCre (truncated form of Cre which is not functional and it is used as control). The next day we replaced the medium with neuronal medium N2/DMEM/F12/NEAA (Invitrogen) containing doxycycline (2 mg/ml, Clontech). We kept the cells in this medium for 5 days, and on day 6 we added ~ 10K mouse glia cells into each 24 well and replaced the culture medium with a serum-containing medium. We analyzed the cultures approximately 3-5 weeks after induction. To generate homozygous knockout neurons, we infected the neurons with LV-Cre or ΔCre one day after induction of the Ngn2 transcription factor.

### Immunofluorescence (IF)

For immunofluorescence (IF) staining of cells (embryonic stem cells and iN cells) we fixed the cells using 4% paraformaldehyde (PFA) for 15 minutes at room temperature and permeabilized the cell membrane using 5% Triton for 1 hour and then blocked the cells in a solution containing 1% BSA, 5% FBS and 1%Triton. The primary antibody was added to the same blocking buffer according to these dilutions: CHD8 antibody (Rabbit-Behtyl lab-A301-224A) used as 1:3,000, Synapsin1 antibody (Rabbit-Synaptic Systems-106002) used as 1:500, Homer1 (Rabbit-Synaptic Systems-160003) used as 1:500, HA (Rabbit-Sigma-H6908) used as 1:500, Map2 (Mouse-Sigma-M9942) used as 1:500,Tuj1 (Rabbit-Biolegened-802001) used as 1:500, ELK1 (Rabbit, Bethyl lab-A303-529A) used as 1:400 and incubated for O/N at 4°C. DAPI added as 100 nM solution for 1 minute. The secondary antibodies were made as 1:1,000 solutions and incubated for 1hr at room temperature.

### Western blotting

Human stem cells and neurons lysed with RIPA lysis buffer supplemented with 5mM EDTA and protease inhibitor (Roche), for 5 minutes at room temperature and 10 minutes on ice. After the lysis, sample buffer (4x Laemmli buffer containing 4% SDS, 10% 2-mecaptaneol, 20% glycerol, 0.004% 4-Bromophenol blue, 0.125 M Tris HCl, pH 6.8) added, and the samples either directly loaded on 4-12% SDS-PAGE gel, or froze in −80 for further analysis. For all of the immunoblots, approximately 20 to 30 μg protein was separated on an SDS-PAGE gel. Antibodies used in this manuscript used with this dilutions: CHD8 antibody (Rabbit-Behtyl lab-A301-224A) used as 1:4,000, ELK1 (Rabbit, Bethyl lab-A303-529A) used as 1:1,000, HA (Rabbit-Sigma-H6908) used as 1:1,000, β-actin antibody (Rabbit,Abcam-ab8227) used as 1:20,000. All blots were visualized by fluorescently labeled secondary antibodies on Odyssey CLx Infrared Imager with Odyssey software (LI-COR Biosciences).

### RNA-sequencing

RNA was obtained from 3 weeks-old cultures of iN cells by adding Trizol LS (Thermo Fisher Scientific) directly into cell culture well. Total 500 ng RNA processed for library preparation using “TruSeq” RNA sample preparation-V2 kit and “Ribo-Zero” rRNA removal kit (Illumina) according to manufacturer’s instruction. The sequencing ran on Illumina’s NextSeq 550 system with 1x 75-bp cycle run.

### RNA-seq data analysis

FastQ files were run on FastQC to obtain high quality (trimmed and cleaned) reads. The reads were aligned to human reference genome sequence (hg19) and assembled with TopHat/Bowtie (version 2.1.1)^33^ for transcriptome analysis. Since we generated the library from a mix of mouse and human RNA, the resulting reads were also from a mixture of both species. We therefore aligned our reads to the human genome with stringent criteria (zero mismatches allowed). The aligned sequences were randomly sampled and re-aligned to the other species’ genome (the mouse, mm9 genome) to ensure that cross-species DNA alignment is not happening. Note that ~ 1% of the reads aligned to both human and mouse genome were discarded from SAM file with SAMtools ^34^. The Refseq hg19 GTF file of transcriptome annotation was downloaded from Ensembl (https://uswest.ensembl.org/index.html) and used as a reference annotation file in TopHat alignment run command to increase the speed and the sensitivity of alignments to splice junctions. Duplicate reads (which arise from PCR step during library preparation) were removed with SAMtools. Pre-built indexes of bowtie were downloaded from the “Bowtie” webpage (http://bowtie-bio.sourceforge.net/tutorial.shtml). AllSAMtools subcommands were used to convert SAM files to BAM files (Bindery Alignment Map). Additionally, SAMtools were used for indexing (to view the signal on genome browser) and for sorting (necessary for downstream analysis). Cufflinks was used for transcript assembly and to estimate the abundance (FPKM) of coding genes. To quantify transcripts across all the samples and obtain estimated counts for downstream analysis, we used HTSeq (htseq-count option) ^35^. These raw counts were used as input for DESeq2 to perform differential expression analysis ^36^ and to generate summarizing plots.

### The single cell RNA data of human brain and the bulk RNA-seq of the developing human cortex

obtained from Allen Human Brain Atlas and the Image credit in figure 1 (with some modification) is the Allen Institute ^6^.

### ChIP-seq and data analysis

ChIP-seq was performed with modifications from a published protocol ^37^. In summary, ten confluent 10cm plates of iN cells (approximately 10×10^6^ neurons in total) 10 days after differentiation was used for chromatin extraction. Cultures were crosslinked with 1% Formaldehyde (Sigma) for 10 min at RT. Glycine (125mM) was added to quench and terminate the cross-linking reaction and after washing with PBS cells were scraped off the dishes and collected into a 50 mL tube. DNA samples were subjected to sonication to obtain an average fragment size of 200 to 600 bp, using Covaris (S220-Focused Ultrasonicator). After sonication, the pellets were cleared from debris by centrifugation in 4°C and the supernatant was collected for further analysis of DNA fragment size (column-purified DNA ran in 2% agarose gel to determine the size) and for DNA/protein concentration analysis. For input calculation approximately 0.5% of cross-linked chromatin separated and saved before the addition of IP antibody. For immunoprecipitation (IP) 1.5 μg anti-CHD8 or anti-HA antibody added into ChIP buffer (RIPA buffer supplemented with protease inhibitors, PMSF and 5mM EDTA) and left to rotate O/N in 4°C. At the same time protein G agarose beads (Active Motif) were washed and blocked with 5% BSA in ChIP buffer and left to rotate O/N in 4°C. The next day, the antibody bound chromatin was added to protein G and rotated 5hr in 4°C. The immunoprecipitated material was washed, and the IP material was eluted from beads with elution buffer (50 mM NaCl, Tris-HCl; pH 7.5) by vortexing at 37 °C for 30min. The eluted DNA was separated from beads with spinning. For reverse cross-linking, the IP and input material was incubated in 65 °C/shaking along with RNaseA (10 ug/ul) and 5M NaCl plus proteinase K (20 ug/ul). DNA was purified on a column (Zymo Research) and processed for library preparation. NEBNext ChIP-seq library prep kit was used for library preparation. Sequencing was performed on Illumina’s NextSeq 550 system with 1x 75-bp cycle run. We obtained 18 to 20 million total reads per sample in one sequencing run.

### ATAC-seq experiment

We followed the Pi-ATAC-seq protocol for the transposition of homozygous knockout and control neurons ^38^. In summary, the cells were fixed in culture for 5 minutes, with 1% PFA and detached from the plate with EDTA and stained for GFP, which allowed us to sort the Cre-GFP positive cells. After that, the transposition proceeded as standard ATAC-seq protocol with slight modification (extra step of reverse cross-linking performed overnight in 65C°). Note that for heterozygous knockout and wild type transposition, we followed the original ATAC-seq protocol in which un-fixed nuclei is permeabilized and subjected to transposition ^39^.

### ATAC-seq, ChIP-seq data analysis

For ChIP-seq and ATAC-seq ENCODE ChIP-seq pipline2 was used to obtain significant peaks^13,18^. For motif discovery, we used HOMER (v4.10) (http://homer.ucsd.edu/homer/). For clustering analysis, we used Cluster 3.0 ^40^. Heatmaps were generated using java program-Treeview ^41^. For ontology analysis, we used DAVID analytical tool ^42^. To obtain estimated counts within the region of interest in ATAC-seq experiment we used FeatureCounts- a general-purpose read count tool from Rsubread package ^43^ and a custom GTF file with the coordinates of the overlapping ATAC-seq peak in all the samples used as input for the program. For library normalization and differential accessibility analysis, we used DESeq2. ^36^. Differential accessible sites (opening and closing regions) were manually examined in UCSC Genome Browser with the 2019 update (http://genome.ucsc.edu). For enrichment analysis and generating normalized heatmaps and signal intensity plots, we used “deepTools” ^44^.

### ChromHMM analysis

We used ChromHMM algorithm to characterize neuronal chromatin state and the functional chromatin domains at CHD8 targets. We obtained histone mark ChIP-seq data of H9 derived neurons from ENCODE portal ^45^. The histone signals binarized across the genome to build a multivariate hidden Markov model and to learn the combinatorial and spatial pattern of histone modification at CHD8 target regions^13^.

### ChIP quantitative PCR (ChIP-qPCR) experiment

Total 5-10 ng ChIP DNA and the input were used to perform a quantitative PCR experiment and measure the enrichment levels. All primers used are listed in “ChIP-seq-peaks.xlsx” file (attached to GSE141085), along with the relevant information, including the closest gene and the number of the motif on the peak. For each peak site, 3 independent technical replicates (independent IP experiments) were used for qPCR analysis. We normalized the ChIP signal over the input signal, which was less than 0.5% for total IP material. Analysis of qPCR experiment performed on the light Cycler 480II (Roche).

### RNA extraction and RT-qPCR experiment for gene expression

For RT-qPCR and RNA-seq experiments, we applied similar RNA isolation methods: neurons that are differentiated on mouse glia cells for ~3 weeks were washed in PBS and then lysed with TRIzol added directly to the plate. RNA was purified with the ZYMO RESEARCH-Direct-zol kit. Human specific primers were used for amplification of desired RNA.

### Analysis of dendritic arborizations

Neuronal cultures fixed at approximately 3 weeks after transgene induction with 4% PFA for 15 minutes. The primary and secondary antibodies dilutions are according to our method in “Immunofluorescence experiment”. For morphological analysis and tracing neurites, we used the MetaMorph ^46^ software and for synaptic puncta analysis and other general image processing, we used java program ImageJand the relevant modules, including CellProfiler 3.0 ^47^.

### AAV-mediated gene targeting

For the generation of conditional CHD8 heterozygous knockout cell line we designed a donor vector for homologous recombination that carries two homology arms around the exon 4 of CHD8 gene and included two loxP sequences in the same direction for frameshifting mutation. A positive selection cassette (neomycin expression to confer resistance to Geneticin) included for purifying clones that carry the integrated donor cassette. The selection cassette contained a splice acceptor (SA) and a sequence for internal ribosomal entry site (IRES) attached to Neomycin resistance gene (NEO) and a polyadenylation (PA) signal. The NEO resistant clones were used for screening PCR to verify the correct inserting of the targeting vector in the locus (see Figure S2-2A to 2C). The PCR primers are designed to cover the region from outside the homology arm (primer # 1 and #4) to inside the cassette.

The drug resistance cassette was flanked with FRT sequence and later removed by transient expression of FlpE recombinase. For HA-FLAG tagging of CHD8 gene, the tags were inserted into the C-terminus region in the frame before the stop codon of Exon 38, together with the Neomycin resistance gene (see Figure S1-1). After infection of ES cells with recombinant AAV (rAAV-DJ) carrying ITR flanked targeting vectors, we selected the cells with Geneticin antibiotic (Gibco) for ten days or until single colonies were obtained. The resistant colonies expanded, and genomic DNA was extracted for downstream analysis.

### Electrophysiology

Electrophysiological recordings in cultured iN cells were performed in the whole-cell configuration as described previously ^7,48^. Patch pipettes were pulled from borosilicate glass capillary tubes (Warner Instruments) using a PC-10 pipette puller (Narishige). The resistance of pipettes filled with intracellular solution varied between 2-4 MOhm. The standard bath solution contained (in mM): 140 NaCl, 5 KCl, 2 CaCl_2_, 2 MgCl_2_, 10 HEPES-NaOH pH 7.4, and 10 glucose; 300-305 mosm/l. Excitatory postsynaptic currents (EPSCs) were pharmacologically isolated with picrotoxin (50 μM) and recorded at −70mV holding potential in voltage-clamp mode with a pipette solution containing (in mM): 135 CsCl, 10 HEPES-CsOH pH 7.2, 5 EGTA, 4 MgATP, 0.3 Na4GTP, and 5 QX-314; 295-300 mosm/l. Evoked EPSCs were triggered by 0.5-ms current (100 μA) injection through a local extracellular electrode (FHC concentric bipolar electrode, Catalogue number CBAEC75) placed 100–150μm from the soma of neurons recorded. The frequency, duration, and magnitude of the extracellular stimulus were controlled with a Model 2100 Isolated Pulse Stimulator (A-M Systems, Inc.) synchronized with the Clampex 9 data acquisition software (Molecular Devices). Spontaneous miniature EPSCs (mEPSCs) were monitored in the presence of tetrodotoxin (TTX, 1 μM). mEPSC events were analyzed with Clampfit 9.02 (Molecular Devices) using the template matching search and a minimum threshold of 5pA, and each event was visually inspected for inclusion or rejection. Intrinsic action potential (AP) firing properties of iN cells were recorded in current-clamp mode using a pipette solution that contained (in mM): 123 K-gluconate, 10 KCl, 7 NaCl, 1 MgCl_2_, 10 HEPES-KOH pH 7.2, 1 EGTA, 0.1 CaCl_2_, 1.5 MgATP, 0.2 Na4GTP and 4 glucose; 295-300 mosm/l. First, minimal currents were introduced to hold membrane potential around −70 mV, next, the increasing amount of currents (from −10 pA to +60 pA, five pA increments) were injected for 1s in a stepwise manner to elicit action potentials. Input resistance (R_in_) was calculated as the slope of the linear fit of the current-voltage plot generated from a series of small subthreshold current injections. To determine whole-cell membrane capacitance, square wave voltage stimulation was used to produce a pair of decaying exponential current transients that were each analyzed using a least-squares fit technique (Clampfit 9.02). Neuronal excitability recordings were performed using standard bath solution supplemented with 20 μM CNQX, 50 μM AP5, and 50 μM PTX to block all possible glutamatergic (AMPAR- and NMDAR-mediated), as well as GABAergic synaptic transmission. Drugs were applied to the bath solutions prior to all recordings. Data were digitized at 10 kHz with a 2 kHz low-pass filter using a Multiclamp 700A amplifier (Molecular Devices). For all electrophysiological experiments, the experimenter was blind to the condition/genotype of the cultures analyzed. All experiments were performed at room temperature.

### Quantifications and statistical analysis

All data are shown as means +-SEM and from a minimum of three biological replicates (independent differentiations). GraphPad Prism and R were used for statistical analysis and calculations of significance.

## Data and code availability

The raw sequencing files are deposited with the Gene Expression Omnibus (NCBI) (GEO accession number: GSE141085). The list of Encode data used in this study listed in “the ChIP-seq-peaks.xlsx” file (attached to GSE141085).

## Notes

### Competing Interest Statement

The authors have declared no competing interest.

